# Feature Design for Protein Interface hotspots using KFC2 and Rosetta

**DOI:** 10.1101/514372

**Authors:** Franziska Seeger, Anna Little, Yang Chen, Tina Woolf, Haiyan Cheng, Julie C. Mitchell

**Author notes:** Correspondence to Julie Mitchell Oak Ridge National Laboratory, Knoxville TN & University of Wisconsin - Madison, Madison WI.

## Abstract

Protein-protein interactions regulate many essential biological processes and play an important role in health and disease. The process of experimentally characterizing protein residues that contribute the most to protein-protein interaction affinity and specificity is laborious. Thus, developing models that accurately characterize hotspots at protein-protein interfaces provides important information about how to inhibit therapeutically relevant protein-protein interactions. During the course of the ICERM WiSDM workshop 2017, we combined the KFC2a protein-protein interaction hotspot prediction features with Rosetta scoring function terms and interface filter metrics. A 2-way and 3-way forward selection strategy was employed to train support vector machine classifiers, as was a reverse feature elimination strategy. From these results, we identified subsets of KFC2a and Rosetta combined features that show improved performance over KFC2a features alone.

## 1 Introduction

Protein-protein interactions play a crucial role in biochemical processes. Modulation of protein-protein interactions bears enormous potential for therapeutic drug development. Thus, accurate predictive models of protein-protein interactions will not only enhance our understanding of the molecular basis of protein recognition and specificity but further provide and inform efforts to modulate protein-protein interactions. Certain hotspot residues at protein-protein interfaces contribute more binding energy to the interaction than others. An *alanine mutagenesis hotspot* in a protein-protein interface is an amino acid for which the change in binding energy upon mutation to alanine exceeds 2 kcal/mol. That is, the change in energy upon binding (*ΔG*_*bind*_) is increased by at least 2 kcal/mol (*ΔΔG*_*bind*_ > 2 kcal/mol). hotspots are known to contribute significantly to the energetics of protein-protein interaction [7, 22, 40]. hotspot analysis has both a long history as well as many recent contributions [1, 2, 4–8, 11, 12, 14–19, 21–25, 28, 29, 33–37, 40, 42, 45–51, 53–57]. Early work on analysis of protein structures in relation to mutagenesis effects established the structural and chemical properties of amino acid residues that significantly alter binding free energy when mutated to alanine [7, 21, 22]. More recent work has begun to characterize hotspot regions, chemical alignment of interfaces, and structural evolution of hotspots [12, 50, 51].

The KFC and KFC2 models for predicting binding interface hotspots [13, 14, 59] have become a gold standard for hotspot prediction. The KFC2 model identifies about 80% of known hotspots [59]. An important recent study of antibody design found the KFC2 model largely in sync with experimental predictions [52]. KFC2 is available via a public web server [14] and has been accessed nearly 80,000 times. The original KFC model examined geometric and biochemical features of a protein-protein interface and used decision trees to develop an accurate predictive model. The KFC2 model pursued a similar line of approach, using support vector machines to train the model and introducing new features that have stronger predictive value than the original ones. In particular, the introduction of interface plasticity measures has significantly improved our ability to distinguish hotspots from non-hotspots.

Rosetta is a molecular modeling and design software suite that has been used for a variety of tasks ranging from protein structure prediction [41] to de novo protein design [26, 30] and protein-protein interface design [9]. Rosetta-based energy calculations [3] have been previously used to create a model for predicting protein-protein interface hotspots [28]. In this work, we will add features from Rosetta to those of KFC2 and train an improved model for protein-protein hotspot prediction. We will combine strategies for feature selection with support vector machine learning in order to achieve an optimal model.

**Fig. 1.**
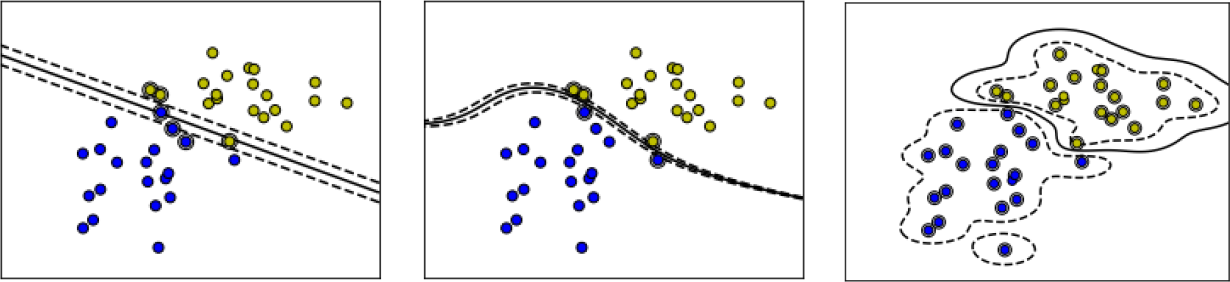
Demonstration of SVM classification with linear (left), polynomial (middle), and RBF (right) kernels. The yellow and blue dots correspond to data points simulated from two multi-variate normal distributions, i.e. two classes. The solid and dashed lines are the contours of the SVM decision function with levels 0 (solid curve) and ±0.5 (dashed curves).

## 2 Background on SVM

Support vector machine (SVM) is a widely adopted binary classifier in recent years due to its efficiency and accuracy [10, 39]. As a supervised classification algorithm, SVM uses labeled training data to build a model and infers the two categories of the testing data. The two categories correspond to hotspot or non-hotspot in our case. In SVM, each data point is represented using a *d*-dimensional vector of descriptors/features and a label that denotes the class (hotspot, non-hotspot). Given labeled training data, SVM identifies a separating hyperplane in the high-dimensional feature space, with each side of the hyperplane corresponding to one (predicted) class. This hyperplane can be used to classify testing data for which the class is unknown. In practice, there are multiple valid hyperplanes that separate the training data. The hyperplane that SVM selects maximizes the distance between the hyperplane and the nearest data point to each side of the hyperplane.

The SVM classifier can be linear or nonlinear, depending on the choice of kernel; see the documentation for SVM from `scikit-learn` [43] for more details. Figure 1 gives an example showing SVM classifiers which use linear, polynomial, and (Gaussian) Radial Basis Function (RBF) kernels, respectively. We choose to use the RBF kernel for our data due to its utility in obtaining the best models for this application. There are two parameters controlling the SVM classifier, *C* which controls the margin between support vectors and the separating hyperplane, and *γ*, which controls the shape of the RBF kernel.

For our training runs, we tabulated performance based on a five-fold cross-validation. Each *C* and *γ* combination is checked using cross-validation, and the combination that leads to the best cross-validation accuracy is selected. See Section 4 for a more detailed description of the SVM implementation and parameter tuning for the hotspot data set.

In the hotspot classification problem, the proportion of hotspots is much smaller than the proportion of non-hotspots. This problem is typically referred to as classification for highly unbalanced data. In this case, the decision function is more driven by the more prevalent class (non-hotspots) instead of the other (hotspots). In order to avoid this issue, we adopt the “class-weighted” SVM: assigning higher misclas-sification penalties to the instances in the rare class and vice versa in the training data so that the decision boundary is almost equally influenced by the two classes. We use the SVC function in `scikit-learn` [43] to implement this.

In interpreting the results of SVM feature selection and parameterization, it is important to understand any resulting model represents *a good choice* rather than *the best choice*. However, we will see some patterns emerge in feature selection if we build a range of models using different parameters.

## 3 Data Sets and Features

The original KFC and KFC2 data sets are described in [13, 14, 59]. For this work, we used a newer expanded data set of alanine mutagenesis hotspots, available from the SKEMPI database [38]. Note that SKEMPI distributes a set of cleaned and renumbered protein structure files that align with their database entries, on which our feature calculations were performed. All KFC2 features were calculated on the structure for the complex, and Rosetta features used relaxed structures and *in silico* mutants.

Structures of the SKEMPI data set of mutant empirical interactions were relaxed in the latest Rosetta full-atom forcefield, REF15, while being constrained to input atomic coordinates [3]. A computational model was generated for each described interface mutation in the SKEMPI dataset by first replacing the native residue with an Alanine residue and performing local side chain minimization within 8 *Å* of the mutated residue. All wild-type and mutant structures were scored with the REF15 Rosetta energy function in addition to seven Rosetta filter terms pertaining to interface characteristics [32]: number of residues participating in the interface, *ΔΔG*_*bind*_ of binding, Larence and Colman interface shape complementarity [31], side chain carbon-carbon contact counts, and a count of the buried unsatisfied hydrogen bond donors and accceptors at the interface. A full description of features is given in Table 1.

We created a custom data set by combining the KFC2a data set with these Rosetta features. Each row in the data set refers to an individual mutation and is labeled as a hotspot or non-hotspot residue based on the empirically determined change in binding free energy [38].

## 4 Feature Selection Strategy and Implementation

The presence of redundant and irrelevant features makes careful feature selection essential, especially for high-dimensional data [58]. We implement a *wrapper* method for feature selection, i.e. the features we select optimize the performance of an SVM classifier. As opposed to *filter* methods, where feature selection is independent of the learning algorithm, wrapper methods treat the learning algorithm as a black box that outputs a performance metric associated with a given set of features, which is then optimized by adjusting the training parameters. This is a simple and powerful approach for feature selection [20].

**Table 1.**
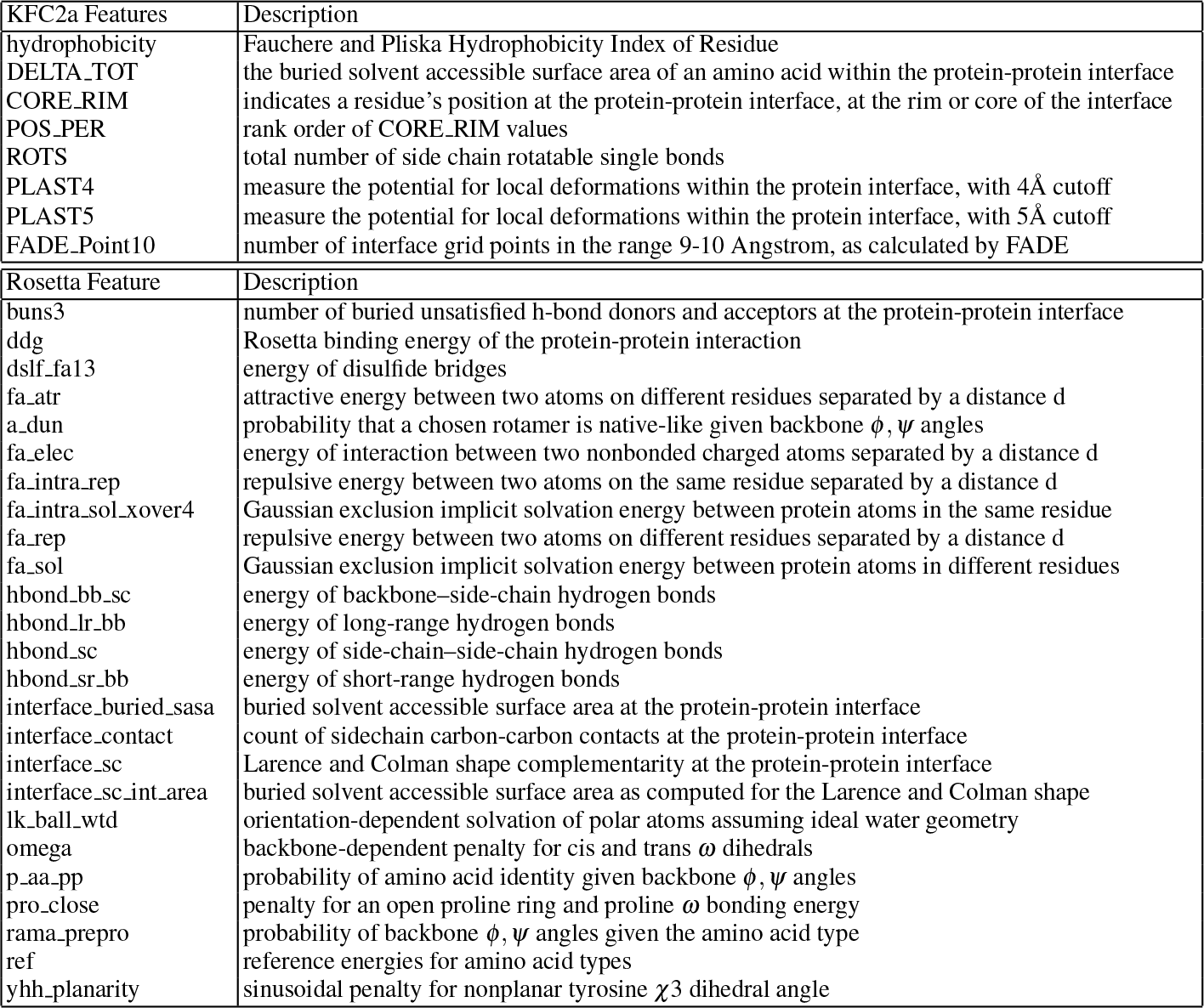
Descriptions of individual KFC2a and Rosetta features used in this study.

More specifically, our goal is to select the set of features that optimizes the cross-validated F1-score of a Gaussian kernel SVM model. The F1-score is a performance metric for a binary classifier, and is defined as:

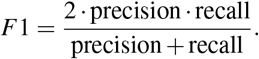

Here, recall is the true positive rate, and precision is the percentage of predicted hotspots that are true hotspots [44]. The F1-score is generally considered to be a more useful measure of performance than overall accuracy, especially when the negative class occurs more frequently than the positive class. To avoid over fitting, we optimize the F1-score using a five-fold cross-validation procedure. Such a validation procedure is intended to mimic the performance on independent test data, by successively eliminating subsets of the data, training on the remainder, and testing on the withheld data. By iterating through folds of withheld data, an unbiased prediction can be made for each data point in the entire data set.

Although we seek the set of features giving us the global optimum of the cross-validated F1-score, it is computationally intractable to search the space of all possible subsets. Assuming we start with *d* features, there are 2^*d*^ possible subsets, giving a complexity exponential in the number of feature combinations [27]. Even if we restrict our search to feature subsets of cardinality *k* ≪ *d*, a brute force search would require that we train *O*(*d*^*k*^) models. For this reason, a greedy algorithm is introduced that selects the highest performing feature and then sequentially grows the feature set; this process is called *forward selection*. This reduces the complexity to *O*(*kd*), though it is easy to construct examples where the feature set obtained is not optimal [20]. In this work we implement an approach which leverages the efficiency of forward selection while reducing the optimization error incurred by the greedy algorithm. We employ a semi-greedy algorithm which at each iteration adds in optimal pairs of features, giving complexity *O*(*kd*^2^). We also compare with the result of adding in optimal triples of features, which has complexity *O*(*kd*^3^). We will refer to the models obtained from forward selection with pairs and triples as Model 1 and Model 2 respectively.

Our SVMs were trained with a custom Python script using the `scikit-learn` library [43]. The features were scaled using the `scikit-learn` preprocessing to have zero mean and unit variance. For every model evaluation, a randomized grid search using a Gaussian RBF kernel and balanced class-weights, distributed to run eight jobs in parallel, was performed to find ideal estimates for the parameters *C* and *γ*. For each parameter combination (*C, γ*) in the random grid, the F1-score was estimated using five-fold cross-validation.

Our feature 2-way and 3-way forward selection strategies are not deterministic due to the computational challenges of fine-grained parameter search, so we ran these parameter searches five times to look for trends among the features discovered, rather than relying on a single run. We also ran Recursive Feature Elimination Cross Validation (RFECV), a reverse selection algorithm for use with linear SVMs for various values of the penalty parameter, *C*. We then trained non-linear classification models using the feature classes identified from the RFECV analysis.

## 5 Results and Discussion

### 5.1 Pairwise Relationships Among Features

First, let’s look at the correlation matrix of features (Figure 2), which are individually described in Table 1. There are two groups of highly correlated features used for KFC2a. The first group (DELTA_TOT, CORE_RIM and POS_PER) are all related in some way to solvent accessibility, and the second group consists of two plasticity features calculated at different distance thresholds (4Å vs. 5Å). Some (mostly weaker) internal correlations exist for Rosetta features. Between KFC2a and Rosetta, the non-trivial correlations were related to buried surface (KFC2a CORE_RIM vs. Rosetta interface_buried_sasa and interface_sc_int_area) and hydrophobicity (KFC2a hydrophobicity vs. Rosetta ref).

**Fig. 2.**
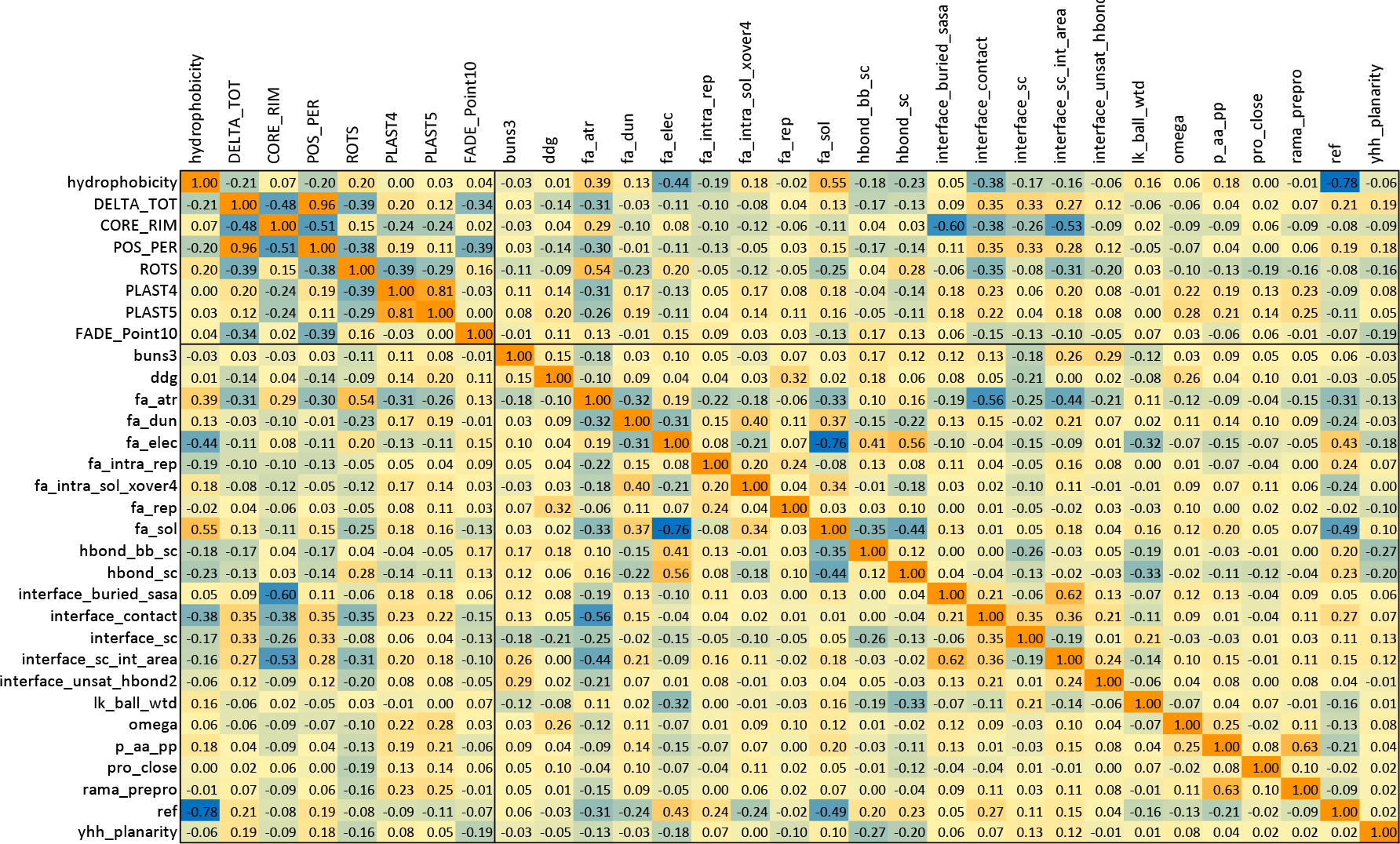
Pairwise correlation scores between features, with significant correlations shown in dark orange or blue.

### 5.2 Recursive Feature Elimination

In addition to our forward selection strategies, we looked at results from the Recursive Feature Elimination Cross Validation method. These models are somewhat easier to analyze, as they are linear models based only on the classification penalty parameter, *C*. Because it is hard to draw conclusions from a single run of the RFE method, we varied the value of *C*, and otherwise used the same SVM training parameters applied in the 2-way and 3-way feature selection strategies.

In Table 2, the feature rank for each feature is shown for varied values of *C*. Notice that the number of features required increases as the penalty for incorrect classification increases, which could lead to a more precise model, but also to over-fitting. Some features such as hydrophobicity appear as a top-ranked feature for every choice of *C*, and we will call this feature group lowC, meaning they can perform well for low values of *C*. In this case, 7/8 KFC2a features and 16/23 Rosetta features are selected. Other features such as fa rep are not highly chosen in any model, and we will refer to such examples as the disfavored feature group. For high *C* values, the RFECV method selects all but a few of the features, and this highC group contains everything but the disfavored features. In Table 2, the lowC features are those in plain text, and the highC features include those in plain and bold text. The features with strikeout text are those in the disfavored group.

**Table 2.**
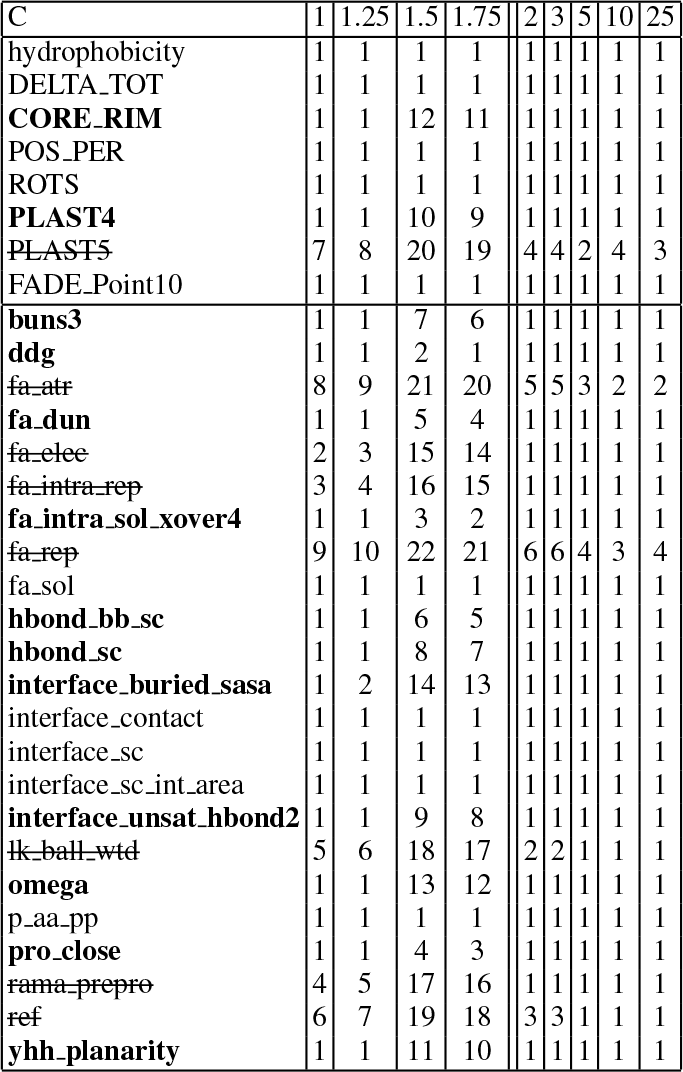
Feature rankings returned from RFECV for each of the 31 features we considered, when examined for various values of *C* between 1.0 and 25.0. The features are grouped into three sets: disfavored features shown in strikeout text, lowC features in regular text, and features in the highC but not the lowC group in bold. The lowC group contains only those features that are top-ranked for all *C* values. The highC group includes everything except the disfavored features.

Using the lowC and highC feature groups, we performed non-linear SVM training using *C* values in the preferred range (0.5 to 2 for lowC; 5 to 25 for highC). We also examined the KFC2a, Rosetta and all features using an exhaustive *C* and *γ* parameter search, using the entire *C* parameter range (0.5 to 25) and finer sampling. These results are shown in Table 3, and we see that the lowC and highC groups return the highest ROC AUC scores when compared with other feature groups. The highC feature group returns the best result overall, with the lowC model performing worse on the positive (hotspot) class. The highC feature group performs similarly to KFC2a features and to the group of all on the positive class (TP+FN) and gives improved predictions on the negative class (TN+FP). The lowC feature group performs similarly to Rosetta features on the negative class, doing somewhat better on the positive class. In Figure 3, we see the results of exhaustive parameter search using non-linear SVMs with radial basis functions on the different feature groups.

**Fig. 3.**
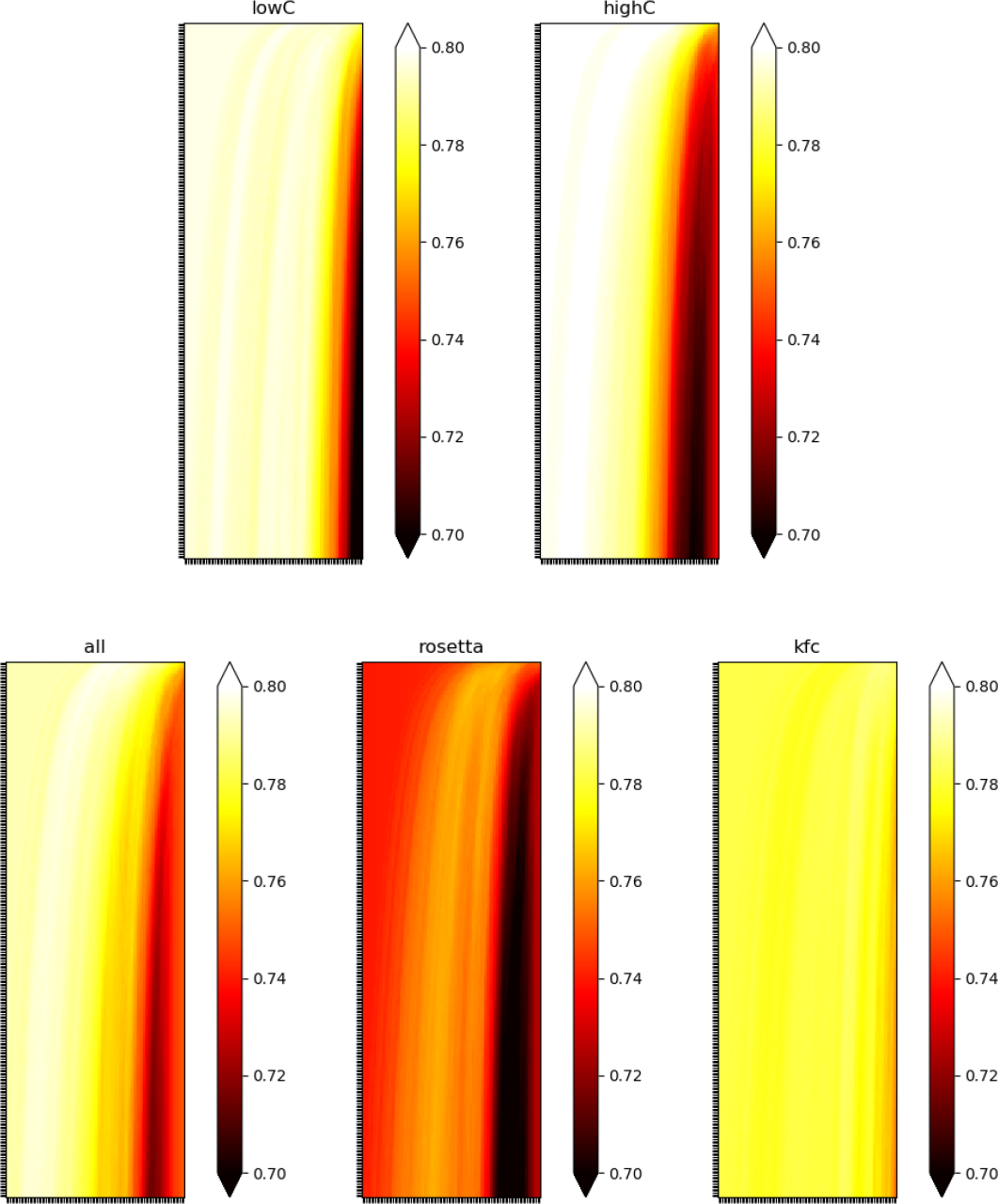
The figure shows ROC AUC scores for models trained on lowC features (top left) and highC features (top right), as a function of *C* and *γ*. The *C* values for lowC vary linearly along the y-axis from 0.5 to 25.0, with 300 samples, and the *γ* values vary logarithmically along the x-axis from 0.00001 to 0.1, with 100 samples. In addition, models trained on all features (bottom left), Rosetta features (bottom middle) and KFC features (bottom right) are shown. The highC feature combination leads to the best chance of finding a high scoring model.

Further examining Table 3, both the lowC and highC feature groups outperform groups consisting of KFC2a features, Rosetta features and all (KFC2a+Rosetta) features. The lowC group performs well on the negative class (TN+FP), while the highC group performs well on the positive class (TP+FN). When comparing the lowC to highC group, the former results in higher precision models and an accuracy similar to the Rosetta feature group, and the latter in higher recall models and specificity similar to KFC2a features or all features combined. The set of all features vs. KFC2a features performs about the same when examined with regard to F1 score and ROC AUC. However, the groups lowC and highC, made by combining KFC2a and Rosetta features, demonstrate improvements in both F1 score and ROC AUC over other feature groups.

**Table 3.** The table shows optimized *C* and *γ* values, the size (N) of each feature group, and confusion matrix entries for cross-validated performance at the optimal *C* and *γ* values for that feature group. In addition, the Accuracy (Ac), Precision (Pr), Recall (Re), Specificity (Sp), F1-score (F1) and ROC AUC are given. The default search range is *C*=(0.5,25) with 300 linear divisions and *γ*=(10^−5^, 10^−1^) with 100 logarithmic divisions; the lowC* and highC* results restrict the *C*-range from 0.5 to 2.0 for lowC* and from 2.0 to 25.0 for highC*. We see from the results that all features are better than simply KFC2a or Rosetta features; however, both the lowC and highC feature groups offer an improvement over all features.

|  | N | C | gamma | TP | TN | FP | FN | Ac | Pr | Re | Sp | F1 | AUC |
| --- | --- | --- | --- | --- | --- | --- | --- | --- | --- | --- | --- | --- | --- |
| KFC2a | 8 | 0.8277591973244147 | 0.03593813663804629000 | 104 | 328 | 150 | 36 | 0.699 | 0.409 | 0.743 | 0.686 | 0.528 | 0.7882 |
| Rosetta | 23 | 1.7290969899665551 | 0.00319926713779738460 | 90 | 362 | 116 | 50 | 0.731 | 0.437 | 0.643 | 0.757 | 0.520 | 0.7657 |
| all | 31 | 0.6638795986622074 | 0.00183073828029536980 | 102 | 334 | 144 | 38 | 0.706 | 0.415 | 0.729 | 0.699 | 0.528 | 0.7991 |
| lowC* | 10 | 0.6010101010101010 | 0.00265608778294668680 | 95 | 359 | 119 | 45 | 0.735 | 0.444 | 0.679 | 0.751 | 0.537 | 0.7990 |
| lowC | 10 | 0.5819397993311037 | 0.00265608778294668680 | 93 | 358 | 120 | 47 | 0.730 | 0.437 | 0.664 | 0.749 | 0.527 | 0.7990 |
| highC* | 23 | 5.6060606060606060 | 0.00031257158496882353 | 105 | 338 | 140 | 35 | 0.717 | 0.429 | 0.750 | 0.707 | 0.545 | 0.8031 |
| highC | 23 | 0.5000000000000000 | 0.00319926713779738460 | 102 | 336 | 142 | 38 | 0.709 | 0.418 | 0.729 | 0.7029 | 0.531 | 0.8060 |

### 5.3 2-way and 3-way Forward Selection of Features

Forward feature selection was performed by adding features in groups of 2 or 3 (2-way or 3-way, respectively) and then selecting the group that best maximizes performance. When doing the forward feature selection, random sampling was used to optimize *C* and *γ* values. Costly searches such as those shown in Figure 3 are not feasible as part of search strategies, but the large regions of good scores suggest random sampling can identify good solutions. To avoid drawing conclusions from a single run of a stochastic method, we ran our forward feature selection five times. In order to compare results with the highC and lowC groups previously discussed, we ran the algorithm with these restricted *C* ranges, in addition to unconstrained random sampling.

It is important to remind the reader of several points about the search algorithm: the parameter search is coarse-grained, random, and based on F1 scores that are not cross-validated. These properties allow the search to run efficiently, but the optimized scores are not directly comparable to cross-validated F1 scores for the trained models described previously. Full parameter searches with cross-validation were conducted with parameters identified using forward selection, in order to compare performance directly with the other strategies, as will be demonstrated below. The Supplementary Materials include the non-cross-validated F1 scores and pairs of features chosen for 2-way feature addition.

For 2-way forward selection, the two features identified in the first iteration, for each of the five runs, were the KFC2a CORE_RIM and POS_PER features. Rosetta’s interface contact score was consistently chosen in second ranked initial pairs, as shown in Table 4.

**Table 4.** For a single run, at the first iteration, the top three results highlight alternative combinations of features that can perform well.

| Pair Rank | non-CV F1 | Feature 1 | Feature 2 |
| --- | --- | --- | --- |
| 1 | 0.53112 | CORE_RIM | POS_PER |
| 2 | 0.52944 | DELTA_TOT | interface_contact |
| 3 | 0.52840 | CORE_RIM | interface_contact |

From the second iteration on, we take the best previous result (in this case CORE_RIM and POS_PER) and search for two additional features to add. KFC2a ROTS is a consistent choice in the second iteration, along with Rosetta omega. At the third iteration, Rosetta’s fa_sol was commonly chosen. Subsequent iterations sample a wide range of KFC2a and Rosetta terms, using them to improve the F1 score. Table 5 shows the feature selection process for four iterations of five runs of the algorithm with *C* sampled between 2 and 25.

**Table 5.**
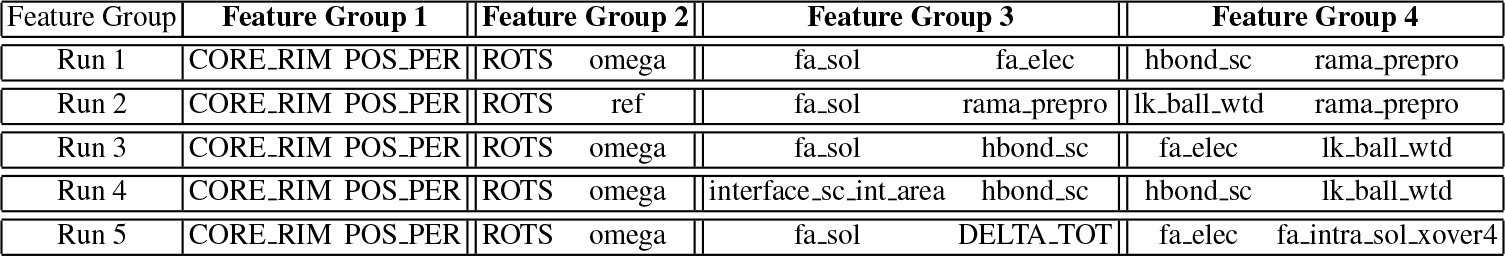
Each group of columns shows the two features added to the model at each iteration, across five runs. As significant improvement in non-cross validated F1 score is observed in iterations 1-2. Around iterations 3-4, the models tend to plateau in performance.

| Feature Group | Feature Group 1 | Feature Group 2 | Feature Group 3 | Feature Group 4 |
| --- | --- | --- | --- | --- |
| Run 1 | CORE_RIM POS_PER | ROTS omega | fa_sol fa_elec | hbond_sc rama_prepro |
| Run 2 | CORE_RIM POS_PER | ROTS ref | fa_sol rama_prepro | lk_ball_wtd rama_prepro |
| Run 3 | CORE_RIM POS_PER | ROTS omega | fa_sol hbond_sc | fa_elec lk_ball_wtd |
| Run 4 | CORE_RIM POS_PER | ROTS omega | interface_sc_int_area hbond_sc | hbond_sc lk_ball_wtd |
| Run 5 | CORE_RIM POS_PER | ROTS omega | fa_sol DELTA_TOT | fa_elec fa_intra_sol_xover4 |

Looking back to Table 2, we see that the KFC2a features POS_PER and ROTS are part of the highC group, and CORE_RIM is in the lowC group. Rosetta’s fa_sol is in the highC group and omega in the lowC group. By the fourth iteration, forward selection has converged. Features like rama prepro and lk_ball_wtd, which were eliminated by reverse selection, begin to appear as selected features but do not offer significant improvements to the model based on F1 scores (see Supplementary Materials.) At this point, feature selection becomes noisy, with many combinations of features offering insignificant improvements to the non-CV F1 score.

The fact that KFC2a features related to core vs rim position of a residue (CORE_RIM, POS_PER) were selected first is a good sign, as core-rim is well known to impact the likelihood of a hotspot. The choice of KFC2a ROTS is not surprising, as it likely reflects some entropic penalty in desolvating long side chains. The choice of Rosetta’s omega is curious, reflecting backbone *ω* angles. However, we see that omega is somewhat correlated to the KFC2a plasticity features, suggesting a correlation for which the cause and effect may be more complex. An unusual omega angle in the mutated structure generated by Rosetta may reflect significant reorganization of the local structure, which is implicitly measured by KFC2a using the plasticity features. In order to compare the forward selection to reverse feature elimination, we trained models using the same parameter search applied to other examples, as displayed in Figure 3. The final ROC AUC for the 8-feature model corresponding to Run 5 of Table 5 was 0.8099, thus exceeding both the lowC and highC feature groups.

**Fig. 4.**
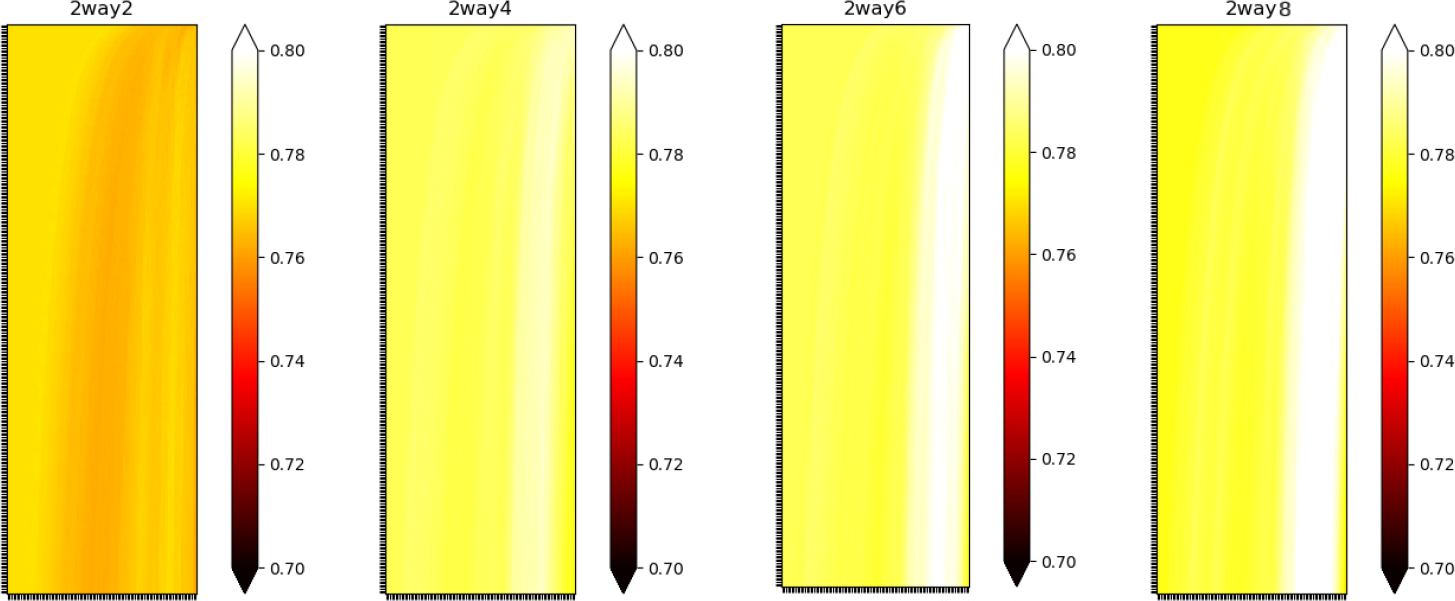
The figure shows ROC AUC scores for models trained on a progression of features identified using 2-way forward selection.

The progression of performance is shown in Table 6, after training the models using cross-validated scoring. In Figure 4, the results of a full parameter search are displayed, using the same bounds used to generate the results of Figure 3. Each iteration increases the zone in which high performing solutions can be obtained. Curiously, the favorable parameter region (high *γ*) that emerges is nearly opposite to that arising from the parameter searches shown in Figure 3. When *γ* is high, the model is localized, and only nearby points influence the prediction at a given instance, whereas when *γ* is small, many points influence the prediction at a single point.

**Table 6.**
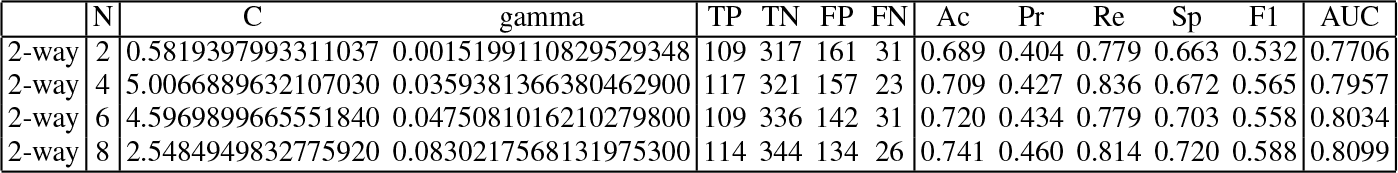
The table repeats the analysis of Table 3 using the results of 2-way forward selection. Results are shown after selecting 2 features, 4 features, 6 features and 8 features. A 5-fold cross validation was used to generate the predictions.

In addition to forward selection adding two features at a time, we performed 3-way forward selection, which showed a very similar progression in selecting features as the 2-way forward selection (Table 7). The 3-way selection showed more variation in initial selection. While CORE_RIM, POS_PER and ROTS was chosen as a good combination, the top combination combined POS_PER with two Rosetta features, interface sc and interface sc int area. Using these three to seed the next iteration, the next three features chosen were DELTA TOT, ROTS and omega, again largely following the preferences of the 2-way search. At the third iteration, buns3, fa elec and interface contact were added. The progression of features selected across five runs of the 3-way forward selection restricting *C* between 2 and 25 are shown in Table 7. The training results and metrics for the 3-way feature selection were fairly comparable to those observed for the 2-way forward selection, hence we omit these details for brevity.

**Table 7.** Each group of columns shows the three features added to the model at each iteration, across 5 runs.

| Feature Group | Feature Group 1 |  |  | Feature Group 2 |  |  | Feature Group 3 |  |  |
| --- | --- | --- | --- | --- | --- | --- | --- | --- | --- |
| Run 1 | POS_PER | interface_sc | interface_sc_int.area | ROTS | DELTA_TOT | omega | buns3 | fa_elec | interface_contact |
| Run 2 | POS_PER | interface_sc | interface_sc_int.area | ROTS | hbond_sc | omega | interface_unsat_hbond2 | fa_dun | hydrophobicity |
| Run 3 | POS_PER | interface_sc | interface_sc_int.area | ROTS | DELTA_TOT | omega | interface_unsat_hbond2 | fa_dun | ref |
| Run 4 | POS_PER | interface_sc | interface_sc_int.area | ROTS | interface_unsat_hbond2 | omega | buns3 | fa_elec | hbond_sc |
| Run 5 | POS_PER | interface_sc | interface_sc_int.area | ROTS | DELTA_TOT | omega | buns3 | fa_elec | interface_contact |

### 5.4 Conclusions

We examined two different strategies for feature selection on a data set for alanine mutagenesis hotspots. The features combined those of a popular hotspot model, KFC2a, and a widely used molecular modeling suite, Rosetta. Recursive feature elimination to define the highC group removed very few features from the combined data set, primarily features that were either redundant or uninformative. The lowC group further reduced the set of features, generally achieving better specificity in prediction than the highC group but lower recall/sensitivity.

An alternate strategy applied forward 2-way and 3-way selection with a random search for optimal *C* and *γ* parameters. These methods converged after just a few iterations, producing a small number of features with significant information content for answering the classification question. The random parameter search was remarkably consistent at finding the top parameter group, CORE_RIM and POS_PER, both of which relate to the “buriedness” of an amino acid within the interface.

The overall preferences for 3-way search versus 2-way search are very similar, but some of the top choices changed. In particular, some Rosetta features that were overshadowed by the dominant choice of CORE_RIM and POS_PER were more prominent in the 3-way search. For example, the CORE_RIM feature, no longer chosen in the initial iteration of 3-way forward selection, is somewhat correlated with both interface sc and interface sc int area, which were chosen instead. This shows the value of considering 3-way forward selection in addition to 2-way selection; in particular, the 3-way selection allowed the first iteration to choose a slightly more accurate combination to cover core-rim effects using three terms.

While showing overall consistency in feature selection, the results also demonstrate that many feature combinations can lead to comparable models. There is not a clearly “right” combination, and the results do not allow us to rank order the importance of any individual feature.

## Supporting information

3way_selection

2way_selection

## Acknowledgements

The feature table and feature selection code are available by email to the corresponding author. We thank the Association for Women in Mathematics (AWM) and the Brown University Institute for Computational and Experimental Research in Mathematics (ICERM) for hosting the Women in Data Science and Mathematics (WiSDM) workshop. The Brown University Center for Computation and Visualization (CCV) and the Institute for Protein Design at the University of Washington provided computational resources used for this project. Participation by JM was sponsored by the National Science Foundation [NSF DMS 1160360]. The AWM Advance Program supported participation by FS, AL, YC, TW and HC. Participation by TW was also supported by DIMACS. FS is generously funded by the Washington Research Foundation Institute for Protein Design Postdoctoral Innovation Fellowship.

